# Assembly of Radically Recoded *E. coli* Genome Segments

**DOI:** 10.1101/070417

**Authors:** Julie E. Norville, Cameron L. Gardner, Eduardo Aponte, Conor K. Camplisson, Alexandra Gonzales, David K. Barclay, Katerina A. Turner, Victoria Longe, Maria Mincheva, Jun Teramoto, Kento Tominaga, Ryota Sugimoto, James E. DiCarlo, Marc Guell, Eriona Hysolli, John Aach, Christopher J. Gregg, Barry L. Wanner, George M. Church

**Affiliations:** Department of Genetics, Harvard Medical School, Boston MA 02115, USA; Wyss Institute for Biologically Inspired Engineering, Harvard University, Cambridge, MA 02138, USA; National Institute of Allergy and Infectious Diseases (NIAID), National Institutes of Health, Bethesda MD 20892, USA; Nuffield Department of Medicine, Medical Sciences Division, University of Oxford, Headington, Oxford OX3 7BN, UK; bioMerieux, Cambridge, MA 02142, USA; Department of Electrical Engineering and Computer Science, Massachusetts Institute of Technology, Cambridge, MA 02143, USA; Sofia University, Sofia 1000, Bulgaria; On leave from Department of Biological Sciences, Purdue University, West Lafayette, IN 47907, USA; Present address: Integrated Research Center of Kobe University, Chuo-ku, Kobe 650-0047, Japan; Department of Biological Sciences, Graduate Schools of Science and Engineering, Tokyo Metropolitan University, Hachioji-shi, Tokyo 192-0397, Japan; Laboratory for Chemistry and Life Science, Institute of Innovative Research, Tokyo Institute of Technology, R1-104, 4259 Nagatsutacho, Midori-ku, Yokohama 226-8503, Japan; College of Physicians and Surgeons, Columbia University, New York City, NY 10032, USA; GreenLight Biosciences, Inc., Medford, MA 02155; Department of Microbiology and Immunobiology, Harvard Medical School, Boston, MA 02115, USA

## Abstract

The large potential of radically recoded organisms (RROs) in medicine and industry depends on improved technologies for efficient assembly and testing of recoded genomes for biosafety and functionality. Here we describe a next generation platform for conjugative assembly genome engineering, termed CAGE 2.0, that enables the scarless integration of large synthetically recoded *E. coli* segments at isogenic and adjacent genomic loci. A stable *tdk* dual selective marker is employed to facilitate cyclical assembly and removal of attachment sites used for targeted segment delivery by sitespecific recombination. Bypassing the need for vector transformation harnesses the multi Mb capacity of CAGE, while minimizing artifacts associated with RecA-mediated homologous recombination. Our method expands the genome engineering toolkit for radical modification across many organisms and recombinase-mediated cassette exchange (RMCE).

## Introduction

Radically recoded organisms (RROs) are desirable for several reasons. For example, recoding enforces genetic and metabolic isolation (*1, 2*), allows facile use of non-standard amino acids (*1, 3*), and has the potential to enable multivirus resistance (*3, 4*). RROs can be used to produce proteins containing non-standard amino acids and are an enabling tool to build biocontained microbes for environmental remediation, industrial applications, and medicine (*5*). In addition, *E. coli* can be engineered to stably maintain non-native sequences from other organisms (*6-9*), support multiplex modification (*10-12*), and deliver DNA to other organisms including diverse bacteria, fungi, plants, and mammalian cells (*13-17*). The ability to transfer scarless large segments of modified DNA to genomes with or without recombination is a desirable feature as genome-scale modifications of organisms across biological kingdoms becomes routine (*9, 12, 18-31*).

In a separate study, we tested 55 of the 87 50-kb segments required to construct a recoded *E. coli* strain in which seven codons were reassigned to synonymous codons, denoted *rE.coli-57* (*5*). The 3.97 MB genome design of *rE.coli-57* has 62,214 codon replacements (*5*). A transformation-based assembly approach allowed us to evaluate each recoded segment individually for strain viability following deletion of the corresponding chromosomal region. To do this, we created a testing pipeline where synonymous codon replacements are validated for strain fitness of each segment (*5*). To construct the fully recoded *rE.coli-57*, we will need to replace all 87 segments throughout the genome at isogenic loci. Previously, we used Conjugative Assembly Genome Engineering (CAGE) to merge sets of genome modifications (*32*) to construct a Genomically Recoded Organism (GRO) where all instances of the TAG stop codon had been replaced with TAA stop codons (*3*). Methods that allow replacement of natural sequences with synthetic DNA segments offer the potential of nearly limitless genome modifications, provided the recoded sequences are viable within living cells (*5*).

When constructing organisms with highly modified genome sequences, there are tradeoffs that will determine which genome assembly technique is optimal. One parameter is how much testing of recoded segments is required to generate optimal phenotypes. The requirement to validate strain fitness favors approaches where both sequences are initially maintained to allow for the systematic deletion of the corresponding natural region to identify deleterious sequences. Direct replacement of segments is supported, however, when assembling strains from prevalidated sequences. A second factor is the optimal size of the replacement segments, which can differ based on the number of mutations or design errors present in synthesized DNA, the need to test DNA segments as they are assembled, and the method used for delivery of the segments by electroporation, phage packaging, conjugation, or transplantation. A final consideration is the requirement for homologous recombination, which can facilitate particular strain assembly methodologies but can also generate undesirable artifacts. *E. coli* offers the ability of using recombinogenic or non-recombinogenic cells, and both the testing protocol (*5*) and CAGE 2.0 can benefit from approaches that minimize recombination.

Although the transformation based method used in our testing pipeline optimizes for the delivery of recoded DNA with minimal opportunity for recombination, the protocol (*5*) is not directly amenable to the assembly of multiple segments. A challenge is the size of the recoded DNA regions that can be delivered. Electroporation can only accommodate sequences of up to several hundred kb (*7*). While it is advantageous to individually test 50 kb segments for effects of recoding on host strain fitness, assembling a genome sequentially from these will be time-consuming. Therefore, a method such as CAGE is ideal because it is amenable to hierarchical assembly (*32*). The testing pipeline also does not easily support the repetitive use of dual selective markers, which are needed to iteratively add additional segments to the same strain at isogenic loci. Techniques described herein will make it possible to extend our testing protocol to larger segment assemblies and, if necessary, to combine the segment assembly and testing protocols.

Here we introduce a next generation protocol, termed CAGE 2.0, for integrating recoded segments into a single *E. coli* strain. In place of the CAGE protocol that uses RecA-mediated recombination for hierarchical assembly, CAGE 2.0 employs λ integrase-mediated site-specific recombination for the targeted assembly of recoded segments. Bypassing the need for the large homology regions that are required for homologous recombination, as well as the need for a recombinogenic background, mitigates against recombination events that can lead to loss of recoded regions during the assembly process. Thus, CAGE 2.0 provides routes to further eliminate undesired recombination, replace segments isogenically, and can be employed to introduce multiple recoded segments in a single strain (Fig. 1).

**Figure 1.**
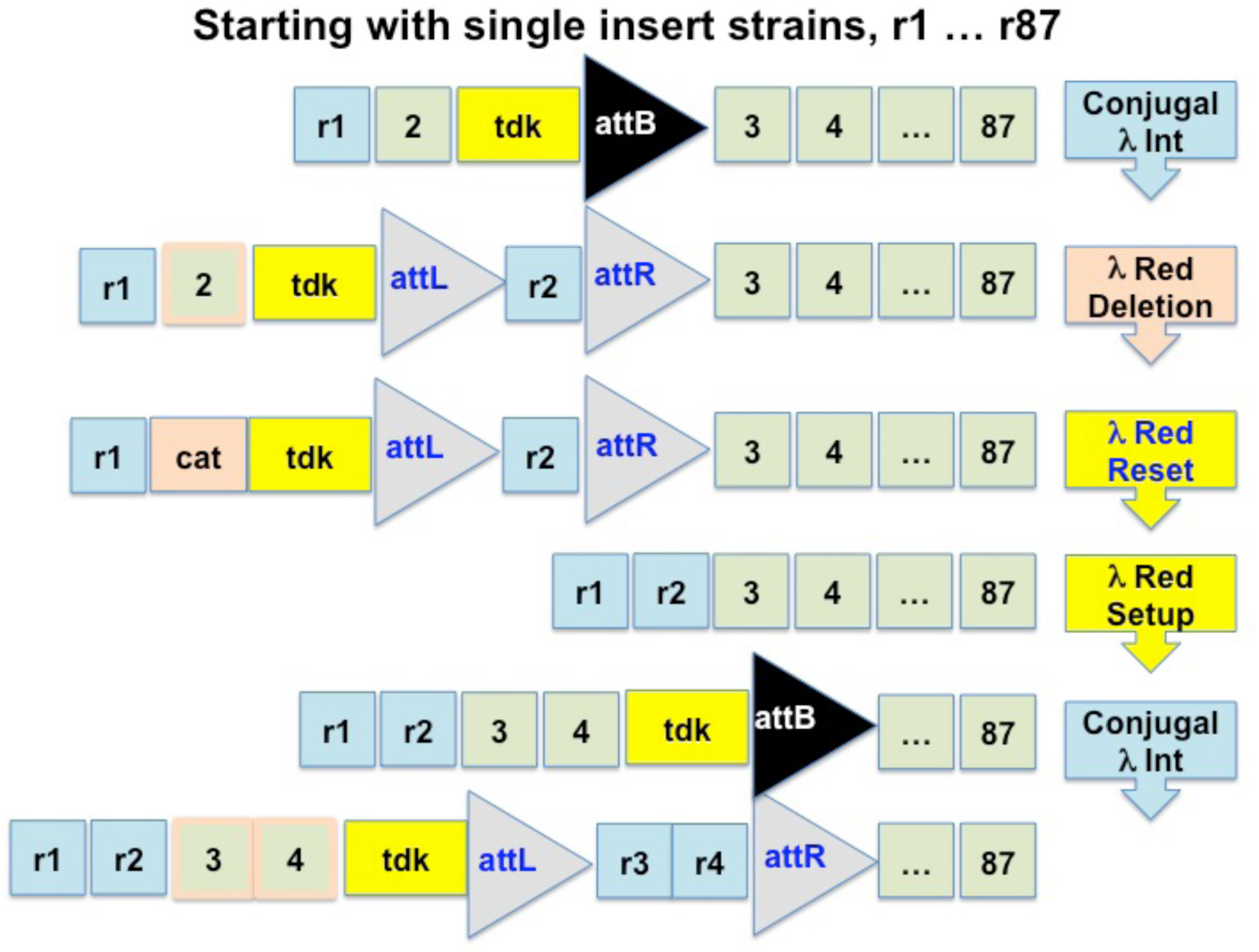
Strategy for CAGE 2.0. **Conjugal λ Integration** of the vector into the recipient strain at single copy. The recipient strain contains the dual selective marker *E. coli* thymidine kinase (*tdk*) and λ integrase *attB* site to the right of the chromosomal sequence that will be replaced. **λ Red Deletion** of chromosomal sequence corresponding to the recoded segment using a chloramphenicol cassette (*cat*). **λ Red Reset Step:** The regions surrounding the recoded segment are removed using *tdk* cassettes and MAGE oligos. **λ Red Setup Step:** To prepare for addition of an additional segment, a cassette containing *tdk* and *attB* is added to the chromosome.

## Results

To expand on the capabilities of CAGE in genome engineering, we devised CAGE 2.0 to deliver recoded segments by site-specific recombination using phage attachment sites that are flexibly shuffled with minimal recombineering to enable genome assembly. Recoded-DNA segments are delivered using vectors that harbor an F plasmid origin, which in *E. coli* has the capacity to replicate extremely large DNA segments ranging from 600 kb to 2.5 Mb (*7, 33, 34*).

In the CAGE 2.0 protocol (Fig. 1), low copy F origin plasmids containing recoded segments are directly conjugated into recipient *E. coli* and site-specifically recombined onto the genome by λ integrase-mediated recombination between the vector *attP* and chromosomal *attB* sites, which are inserted at defined loci by recombineering. Next, the corresponding chromosomal segment is eliminated by λ Red-directed exchange using a chloramphenicol resistance cassette. The resultant strain is then modified for subsequent use as a recipient by removal of residual vector sequences and *att* sites flanking the recoded segment in sequential steps using MAGE (*10*) and *E. coli* thymidine kinase *(tdk)* as a dual selectable marker (*35*). Finally, the host is set for introduction of the next segment by introduction of a new *tdk-attB* cassette in the locus where the recoded segment is to be integrated (Fig. 1). A novelty of the protocol is the use of *tdk*, a marker that can be used repeatedly in the same strain to assist in either adding or removing elements from the genome, allowing for segment assembly.

While developing the testing protocol, we initially attempted to integrate recoded segments both close to and far away from their natural location in *E. coli* K-12 BW38028 (*36*). In this recombinogenic (RecA^+^) strain, we found recombination between natural and recoded segments sequences was problematic at the isogenic locus. Accordingly, we chose two precautions in our initial CAGE 2.0 experiment. We elected to adjacently assemble two recoded segments (segments 43R and 47R (*5*)) in genomic locations greater than 1-Mb away from their natural locations in order to minimize recombination. We also used the *recA1* host *E. coli* K-12 DH10B-MAGE for subsequent CAGE 2.0 experiments to minimize recombination (*5, 37, 38*). By expressing λ integrase during the conjugation process, we were able to recover single-copy recoded segment integrants from the resulting population of cells. Proceeding directly with these clones, we reset the strain by eliminating residual vector sequences and the *attL* and *attR* sites created by integration (Fig. 1).

Performing this process repeatedly and efficiently required the use of a robust dual selectable marker. Our tests with *E. coli tdk* showed that this alternative marker allowed for the maintenance of a stable selective advantage (analogous to *tolC* (*39*)) over multiple rounds of selection using commercially available reagents in a single strain (Fig. 2A, B, and C). Using positive and negative selection conditions optimized for use in GROs (*1*), we found that *E. coli tdk* was more robust than herpes simplex virus thymidine kinase (*hsv-tk*) (*40*) over multiple rounds of selection. When optimizing the negative selection for *tdk*, we circumvented the use of dP because it is a potent mutagenic reagent (*40*), and instead used AZT, a chain-terminator that results in failure of cell division at high doses (*41*) (Fig. 2A and B; see the Supplement for details.) *E. coli tdk* is also attractive because it can be used in a wide range of organisms (*42, 43*), and can maintain utility over multiple rounds without requiring extensive strain modifications, as was required to optimize for *tolc* selection (39).

**Figure 2.**
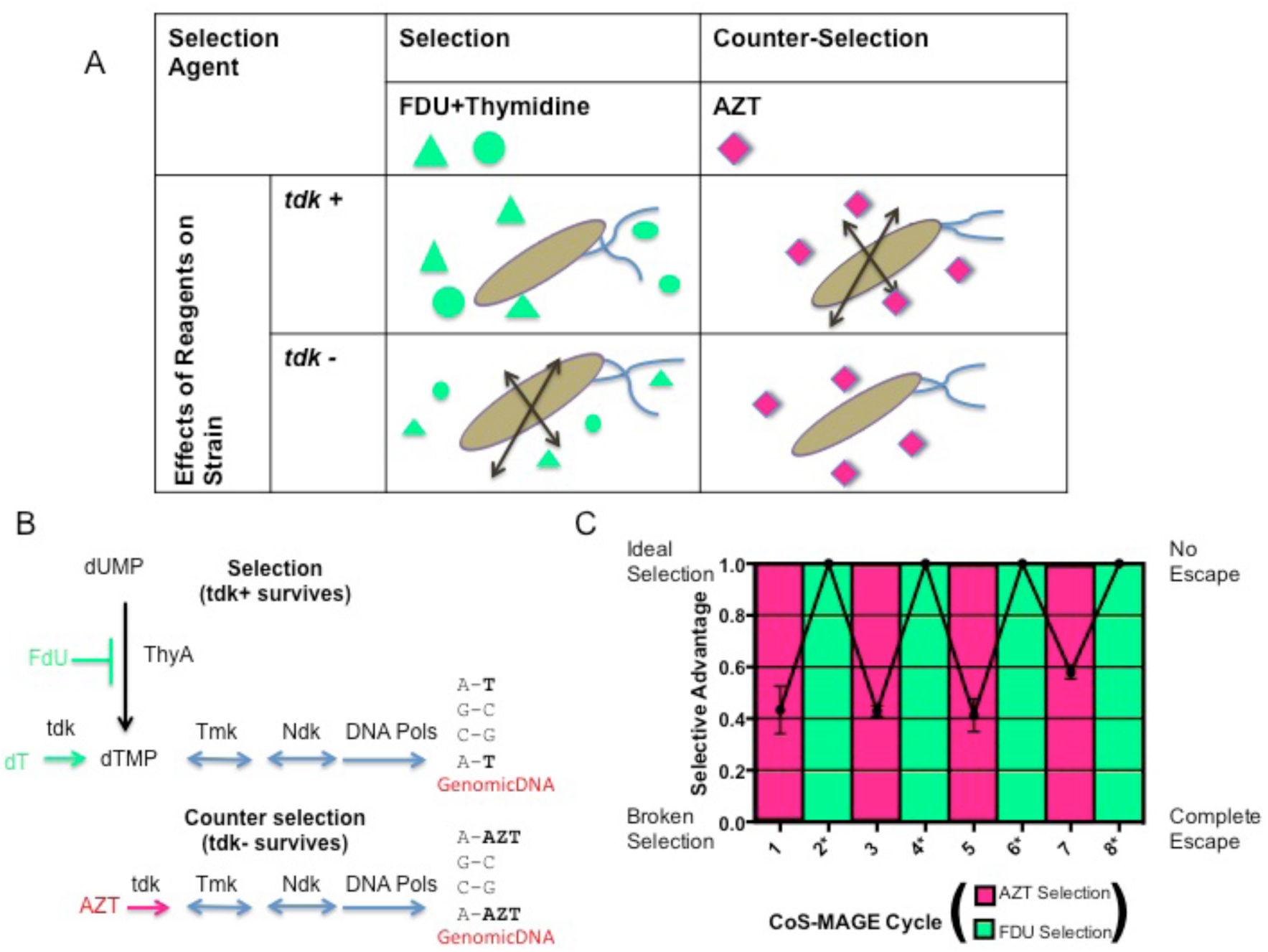
Repetitive *tdk* selection and counterselection maintains a stable selective advantage over multiple cycles (A, B, and C). A) Using a selection reagent FDU utilized for herpes simplex virus thymidine kinase (*HSV-tk*) (*40*), but replacing the mutagenic counterselection reagent dP (6-(β-D-2-deoxyribofuranosyl)-3,4-dihydro-8*H*pyramido[4,5-c][1,2]oxazin-7-one) with the chain terminator AZT, we developed selection and counterselection conditions for *tdk* using commercially available reagents. B) Selection Mechanism of Action: *Tdk* rescues strains in the presence of FDU, which blocks thymidine salvage. Counterselection Mechanism of Action: The presence of *tdk* allows AZT, a chain terminator that inhibits DNA synthesis, to be incorporated into DNA after phosphorylation with *tdk* (*47, 48*). C) To test the robustness of reuse of the *E. coli tdk* gene over multiple cycles in the same strain, we tested the maintenance of the strain to selective advantage during co-transformation with a set of MAGE oligos designed to implement modifications over 10 genomic locations.

By using *tdk*, we developed a protocol to reset the strain after addition of the recoded segment 47R (*5*) (Fig. 1). Initially, we experienced difficulty eliminating the integrated pYES1L vector sequences containing the F replication origin. By using MAGE and negative selection of the adjacent *tdk* marker, we were able to delete the plasmid backbone. The region containing the other attachment site was easily removed by swapping with *tdk* and introduction of a new *attB* site. We subsequently integrated segment 43R at the new *attB* site adjacent to 47R.

When we tested growth of the strain containing the adjacent 43R and 47R segments, we found the normalized generation time was significantly slower than strains containing either sequence alone (Fig. 3A and B). This result motivated us to consider the importance of context in strain development and re-examine alternative routes to assembly of the recoded segments. As a test case, we chose segment 16R (*5*) for isogenic replacement in a *recA1* host. Using the CAGE 2.0 protocol, we first integrated segment 16R on the genome adjacent to the natural locus. Attempts to eliminate the corresponding wild type region together with *tdk* and *attL* using MAGE and negative selection resulted in strains exhibiting recombination between the natural and recoded segment. As an alternative, we first removed the wild type segment by λ Red-directed exchange for a chloramphenicol resistance cassette and then deleted *tdk* together with vector and *attL* sequences using MAGE. Subsequent removal of *attR* site with *tdk* selection enabled the isogenic replacement to be completed.

**Figure 3.**
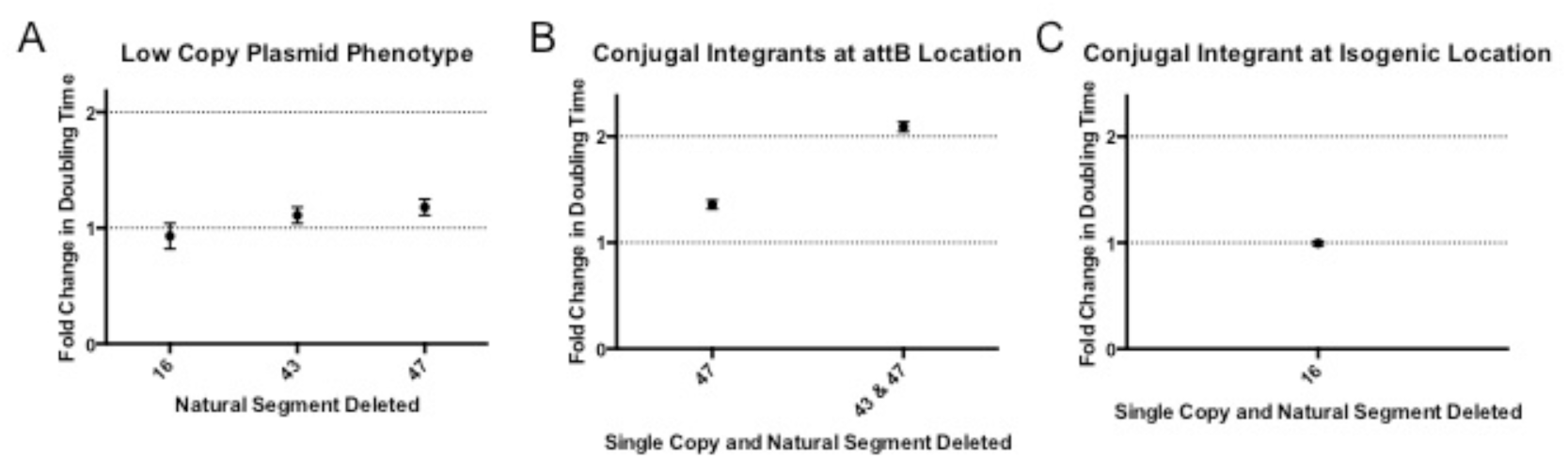
Phenotypic analysis of adjacent and isogenic recoded segments. A) Phenotype established by transformation of recoded segment vector and deletion of the corresponding wild type region (*5*). B) Phenotype established by iterative assembly of two segments at a non-natural location. C) Phenotype established for an isogenic segment replacement.

Scarless replacements are potentially important for preserving regulatory features in adjacent loci that could affect the expression or function of genes within recoded segments that are required to maintain strain fitness. Analysis of the growth curve of the segment 16R isogenic replacement revealed a growth rate similar to the parental wildtype strain, suggesting isogenic replacements that preserve the genome context of the segment are useful intermediates for subsequent assemblies (Fig. 3C). The integration of recoded segments at natural loci is a key consideration for optimizing strain phenotypes and will eventually be a prerequisite for fully isogenic construction of *rE.coli-57*.

## Discussion

The generation of RROs on practical timescales requires improved technologies for assembling modified genomes from large synthetic DNA sequences. These methods would ideally offer the ability to easily transfer large segments of recoded DNA into host organisms and efficiently integrate them into the genome at their corresponding locus. A primary obstacle with many current techniques in *E. coli* is the occurrence of genetic recombination between wild type and recoded sequences after delivery into host strains. As we demonstrated in our testing pipeline, one way to circumvent this problem is to retain the recoded segment episomally in a *recA1* strain, while deleting the corresponding segment from the host chromosome prior to integration. Combining this order of operations with CAGE 2.0 may provide a facile route to generate isogenic replacements. As shown previously, this approach requires an additional Cas9 step to eliminate residual free delivery vector following integration of the recoded segment (*5*) and protocols to subsequently remove the Cas9 plasmid (see Supplemental Methods).

As we move towards generation of *rE.coli-57*, hierarchical methods of assembly will dramatically reduce the time to final strain construction compared to sequential addition of 87 segments. For hierarchical assembly using CAGE 2.0, the protocol must support the excision, maintenance, transfer, and integration of increasingly large recoded genome segments. Currently, genomic excision experiments support use of this protocol for at least 8 segments (see Supplemental Methods), indicating this approach has the potential to drastically reduce the number of steps required for hierarchical assembly.

An alternative to the use of the large vectors required for CAGE 2.0 would be the development of a conjugation-based RMCE method, where very large recoded genome regions flanked by orthogonal *attP* sites are transferred from donor to recipient strains and integrated at isogenic loci. This method takes advantage of well-characterized attachment sites commercially utilized in Gateway products, which are capable of unidirectional integration. With CAGE 2.0, many tools for the task have been prototyped, such as *tdk*-mediated insertion and removal of recombinase sites, integration that is simultaneous with conjugation, and strong markers that select for the delivery of conjugated DNA into the recipient strain. A potential hurdle to overcome in developing this combined approach is the possibility of undesired recombination observed previously using RMCE in recombinogenic (*recA*^+^) *E. coli* (*28*), and which potentially can occur due to reversions from *recA1* to *recA*^+^. An ideal host strain for RMCE + CAGE 2.0 would exhibit a low frequency of unwanted recombination, favoring predictable and selectable isogenic replacements.

We have shown that CAGE 2.0 in *E. coli* can be facilitated by using *recA1* host strains that reduce undesired recombination events. Performing this protocol in other organisms where recombination is essential or where each modification is time consuming will provide a greater challenge. In these cases, achieving isogenic replacements will likely require deletion of the corresponding wild type locus before recoded segments are integrated. If double crossovers can be successfully minimized, then backgrounds that enable recombination could offer advantages for organisms that are normally resilient to modification. A potential approach would be to partially delete the wild type sequences (before segment integration), leaving the flanking homology arms intact, allowing for subsequent integration screening using *tdk* or fluorescent marker expression. In this case, the presence of cellular recombination pathways may offer an endogenous route to strain reset. This would be desirable in hard to modify strains such as mammalian cells, where advance deletion of one chromosomal fragment using CRISPR (*44, 45*) (i.e., before the new sequence in introduced into the cell) may not be disruptive to the cell (due to the presence of multiple chromosomes) and where bacterial artificial chromosome (BAC) libraries provide an engineerable substrate that contains large homologous regions.

As one begins to modify BACs for genome project write (GP-Write) (*31*) and deliver them to mammalian cells or other organisms, the CAGE 2.0 protocol may be directly useful in a number of ways. Conjugal integration provides a route to reduce BAC vectors previously utilized for genome sequencing to single copy and proceed with either MAGE modification or replacement of synthesized regions assisted by positive and negative *tdk* selections, within a strain suitable for BAC maintenance (*38*). After modification, these vectors can be excised or potentially transferred to other strains as genome integrants using RP4 conjugation (*15*). If necessary, several segments can be assembled together using the same protocol prior to transfer to another strain. The ability to modify and transfer BAC vectors and genomes into mammalian cells and other organisms from biocontained *E. coli* (*1,3*) is even more advantageous, offering a method that maximizes biosafety.

The CAGE 2.0 protocol described here further extends the technologies available for constructing RROs. Our results highlight potential opportunities and challenges for each in recoded genome assembly (Fig. 4). Both for these protocols and as one moves towards conjugation and integrase assisted genome engineering in *E. coli*, it will be desirable to further minimize recombination proficiency, which may allow for automation of the protocol. Recently, we have utilized Cas9 to eliminate strains that exhibit undesired mutations, which could assist in minimizing *recA1* reversion (*46*). With CAGE 2.0, we have generated a new set of tools that expands current capabilities in genome construction, and are now developing RMCE + CAGE 2.0 to facilitate large-scale hierarchical approaches to RRO assembly.

**Figure 4.**
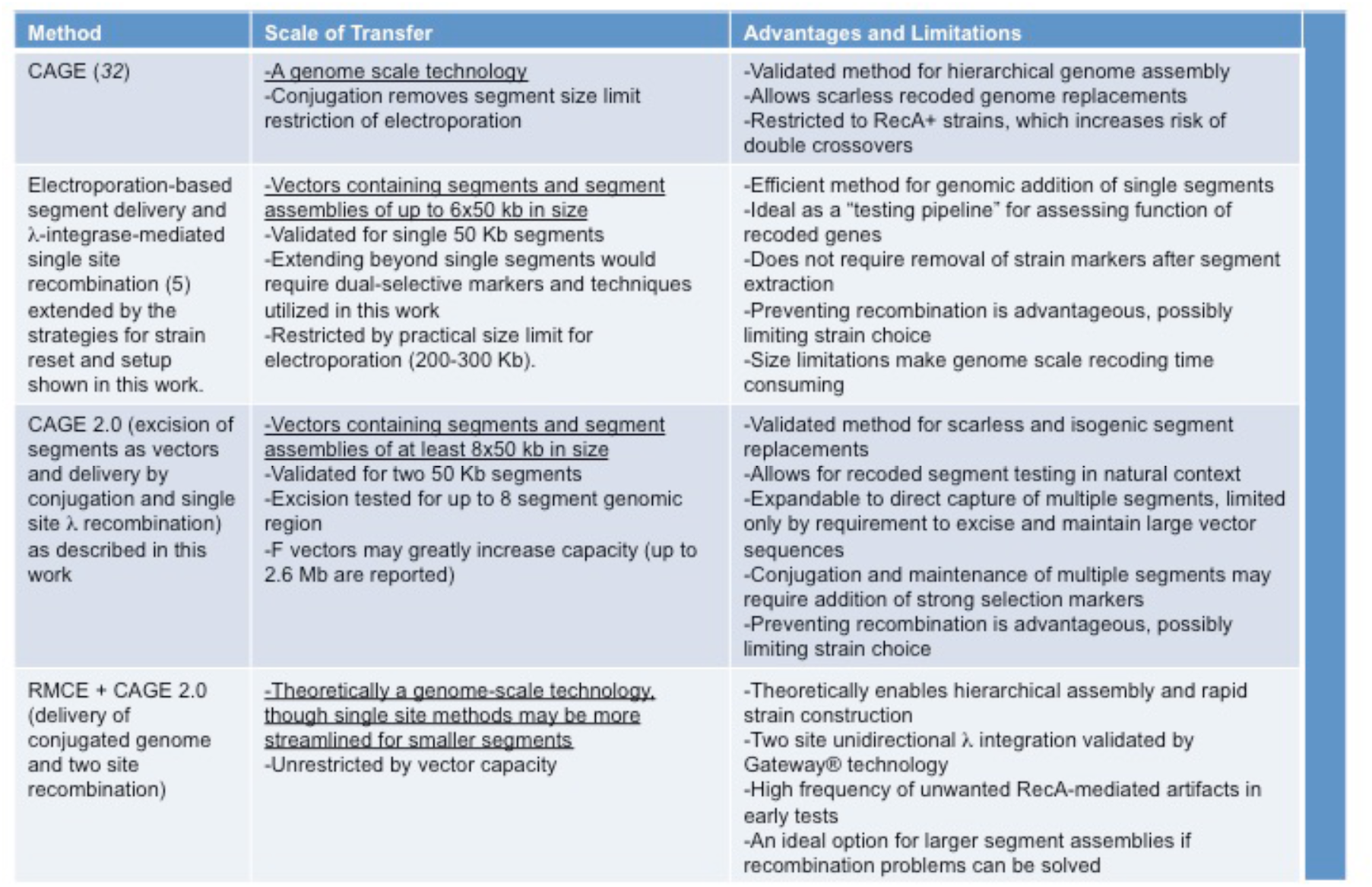
Survey of methodologies for assembling radically recoded genomes using 50 kb segments.

## Acknowledgments

This research was supported by Department of Energy grant DE-FG02-02ER63445 to G.M.C., Synthetic Biology Engineering Research Center from the National Science Foundation SA5283-11210 to G.M.C., DARPA contract HR0011-13-1-0002 to G.M.C., NSF grant number 106394 to B.L.W., Summer Honors Undergraduate Research Program Grant to E.A., and Origins Summer Undergraduate Research Grant to M.M. We thank Timothy Henion for writing assistance on this manuscript

